# Colonization of phosphate-solubilizing *Pseudomonas* sp. strain P34-L in the wheat rhizosphere and its effects on wheat growth and the expression of phosphate transporter gene *TaPT4* in wheat

**DOI:** 10.1101/294736

**Authors:** Xixi Liu, Xiaoxun Jiang, Weirong Zhao, Yuanyuan Cao, Tingting Guo, Xiangyi He, Haiting Ni, Xinyun Tang

**Author notes:** Present address: Hefei Product Quality Supervision and Inspection Institute, Hefei 230088 Anhui Province, P. R. China. X.L. and X.J. contributed equally to this work.

## Abstract

The ability to colonize the rhizosphere is an important basics requirement for field application of plant growth-promoting rhizobacteria (PGPR) strains. There are complex signal exchanges and mutual recognition between microbes and plants. In this study, phosphate-solubilizing *Pseudomonas* sp. P34, a PGPR strain with affinity to wheat, was isolated from the wheat rhizosphere by wheat germ agglutinin (WGA). The plasmid pTR102 harboring the luciferase *lux*AB gene was transferred into P34 to create P34-L. The labeled strain was used to track the temporal and spatial characteristics of colonization in wheat rhizosphere and its effects on wheat development. The transcript level of phosphate transporter gene *TaPT4*, a phosphorus deficiency indicator gene, in wheat roots was monitored by quantitative reverse-transcription PCR. The experimental results indicated that there was a high density of stain P34-L within the top 8-cm depth of the wheat rhizosphere on day 36 of wheat growth. The strain could survive in the wheat rhizosphere for a long time, and colonize new spaces in wheat rhizosphere following the extension of wheat roots. Compared with uninoculated wheat plants, those inoculated with P34-L showed significantly increased phosphorus accumulation in leaves, seedling fresh and dry weight, root fresh and dry weight, total root length, and number of root tips, forks, crossings, which showed a great value of application of the strain on wheat production by promoting the root growth and dry matter accumulation. Strain P34-L down-regulated the transcript level of *TaPT4* in wheat roots, which means a well phosphorus supplementation environment was established by P34-L.

**Importance:** Many PGPR strains often failed to achieve the desired effects when applied in the field. One major reason for the failure is lack of a special affinity between a certain strain and the target host plant, so those strains have low competitive ability with the indigenous microorganism, and unable to survive constantly in rhizosphere. In this work, a new technique to isolate wheat-specific phosphate-solubilizing PGPR strain by WGA was established. The isolate P34 was confirmed can colonize the wheat rhizosphere, and have significantly ability in promoting phosphorus absorption and wheat growth by luminescence labeling techniques. Furthermore, the phosphate-solubilizing ability of this affinity PGPR strain was verified in gene level by quantitative reverse-transcription PCR. These results lay a firm foundation for further research on the relationships between PGPR and their host plants. Meanwhile, this work supplied a potential ideal biofertilizer producing strain for sustainable agriculture.

## Introduction

The rhizosphere, which is defined as the soil space affected by plant roots, plays an important role in plant growth and nutrition (1, 2). Plants, soil, microbes, and the environment constitute the rhizosphere ecosystem. Matter, energy, and signals are exchanged frequently in this ecosystem. Plants and microorganisms interact and stimulate each other’s metabolism, growth, morphology, and development. This forms a stable rhizosphere ecosystem, and ensures plant growth and nutrition (3, 4).

The Plant growth-promoting rhizobacteria (PGPR) are a heterogeneous group of beneficial bacteria that inhabit the plant rhizosphere. They can improve plant growth, prevent disease, and increase yield. They directly or indirectly promote plant growth by fixing atmospheric nitrogen, dissolving phosphorus, releasing potassium, producing phytohormones, and secreting antibiotics (5). There is more than 30 years of research on PGPR (6), and most studies have focused on the selection and application of excellent PGPR strains (7). However, due to a lack of reliable data and technology, there is no guarantee that there will be a special affinity between a certain strain and the target host plant. Therefore, it becomes one of the main reasons for fluctuanting effects in application of PGPR, especially in field conditions (8, 9).

The long-term and stable colonization of PGPR in the rhizosphere of the host plant is a key step for a constant PGPR–host relationship. There are complex signal exchanges and mutual recognition between microbes and plants. Lectins play an important role in recognition in hosts and microbes. Lectins are produced by the host and can reversibly and nonenzymatically bind to specific carbohydrate groups located both on the surface of plant roots and on the surface of bacteria. This initiates a reaction that leads to the successful colonization of the rhizosphere by bacteria (10–13).

In spite of the importance of PGPR for plant growth, few studies have focused on the crucial steps of how these microbes stably colonize the plant rhizosphere. Investigation of survival and colonization of host plant rhizosphere by PGPR require suitable detection methods to trace the target microbes in the environment containing a large number of homologous indigenous bacteria (14). The establishment and development of labeling systems provides effective means for the study of survival status and population numbers of these strains in the plant rhizosphere (15). Among these labeling systems, luminescence labeling techniques have been applied extensively for its high specificity and convenient in many fields, such as tracking pathogenic bacteria in plants, detection rhizobia in legume nodules, monitoring bacteria in the environment and studies on microbial ecotoxicology (14, 16, 17). As a luciferase structural gene, *lux*AB gene is a promising marker for detection of marked strains in the rhizosphere. For luminescent colonies of *lux*AB-marked strains can just be observed by the unaided eye in the dark, it is possible to determine and count the target bacteria simply with dilution plate counting method (18). Use of *lux*AB luminescence labeling techniques has enabled detection of luminescent bacteria even when they were substantially outnumbered by the indigenous microorganisms (15).

Plant root exudates affect the composition and distribution of microbial flora in the rhizosphere. At the same time, the activities of microorganisms in the rhizosphere influence and regulate plant growth and nutrient absorption. This process may be carried on by physiological responses of the host plant to microbial action. To date, most studies on this field have focused on legume-rhizobia, plant-mycorrhizae symbiotic relationships and plant-pathogenic microorganisms parasitic relation (19–22), but few studies have concentrate on beneficial microorganisms of non-leguminous plants, especially graminaceous crops.

For plant root system, the processes of phosphorus absorption from environment and transfer of the phosphorus nutrient to tissues and cells are mediated by a phosphate transporter (PT) located in the plasma membrane (23). There have been several reports on phosphate transporter genes in wheat (24–26), but most studies on this field have aimed only at function of these genes and regulation of phosphate transporters by arbuscular mycorrhizal fungi in wheat (24, 26, 27–29), and few studies have focused on regulatory effects of PGPR on these genes in wheat. *TaPT4*, one member of high-affinity phosphate transporter gene, was found to have strong affinity to phosphorus, and to play a significant role in the absorption of phosphorus under conditions of low phosphorus stress in wheat (26,30–31). Studies indicated that phosphorus supplementation can down-regulate the expression of *TaPT4 gene* in wheat roots (26). So the *TaPT4* gene could be used as a phosphorus deficiency indicator gene to reveal the ambient phosphate level. In this work, a phosphate-solubilizing rhizobacterium *Pseudomonas* sp. strain P34 with affinity to wheat was obtained from the wheat rhizosphere by a screening method of phosphate solubilizing selective medium combined with wheat germ agglutinin (WGA). To track the colonization dynamics of the target strain in the wheat rhizosphere, the luciferase *lux*AB gene was introduced into this strain by electrotransformation to create P34-L. We investigated the colonization dynamics of the phosphate-solubilizing rhizobacterium, *Pseudomonas* sp. strain P34-L (harboring the *lux*AB gene), and analyzed its growth-promoting effects on wheat. We also investigated the effects of this rhizobacterium on the transcript levels of a phosphate transporter gene *TaPT4* for preliminary exploration the molecular mechanisms of its promoting activity on phosphorus absorption for wheat.

## Materials and methods

### Plant materials, strain, media, and soil

Wheat (*Triticum aestivum*W52) seeds were obtained from the School of Agronomy, Anhui Agricultural University, Hefei, Anhui, China. *Escherichia coli* WA803 harboring the plasmid pTR102, which contained the *luxAB* gene and tetracycline and kanamycin double resistance markers, was supplied by College of Life Sciences, Nanjing Agricultural University, Nanjing, Jiangsu, China.

The phosphate growth medium (32) contained the following constituents (g⋅L^−1^): glucose10.0, Ca_3_(PO_4_)_2_ 5.0, MgCl_2_⋅6H_2_O 5.0, MgSO_4_⋅7H_2_O 0.25, KCl 0.2, (NH_4_)_2_SO_4_, 0.1, pH 7.0. The LB medium contained the following constituents (g⋅L^−1^): Bacto-tryptone 10.0, yeast extract 5.0, NaCl 10.0, pH 7.0. The SOC medium contained the following constituents (g⋅L^−1^): peptone 20.0, yeast extract 5.0, NaCl 0.5, KCl 0.186, MgCl_2_⋅ 6H_2_O 0.95, MgSO_4_⋅7H_2_O 1.2, glucose 3.6, pH 7.0.

The yellow cinnamon soil used in this study was collected from Anhui Agricultural University, and was sieved through a 2-mm mesh. The basic physico-chemical characteristics of the soil (g⋅kg^−1^): organic matter 19.62, total N 1.16, available P 0.0206, available K 0.138, pH 6.55.

### Isolation and identification of phosphate-solubilizing bacterial strains

Phosphate-solubilizing bacterial strains were isolated from wheat rhizosphere soil in the city of Hefei (117.27 ° E; 31.85° N) in Anhui province, China. The wheat (*Triticum aestivum* W52) plants were uprooted carefully and removed the excess soil particles by vigorously shaking. One g of rhizosphere soil adhering to the wheat root surface was transferred to a 250 ml Erlenmeyer flask with 100 ml of sterile water in it. After shaking for 30 min at 150 rpm in shaker, serial dilutions of rhizosphere soil were prepared, and 0.1ml of each 10^−4^ to 10^−6^ dilutions were spread on phosphate growth medium plates. Phosphate-solubilizing bacteria were identified based on an obvious halo around their colonies after 5 days of incubation at 28°C. Pure cultures were isolated by streaking repeatedly on plates containing the same medium.

Phosphate-solubilizing bacteria with affinity to wheat were rescreened by WGA. WGA was purified by precipitation with ammonium sulfate and affinity chromatography from wheat seeds as described by Levine *et al* (33), then diluted with 10 mM⋅L^−1^phosphate-buffered saline (pH 7.2) to 2 mg⋅mL^−1^, and stored at 4°C.

The isolates were cultured in LB medium for 15 h at 28°C with shaking (160 r⋅min^−1^). The cells were collected by centrifugation at 4000 r/min for 8 min, and washed three times in sterile distilled water, and then re-suspended in sterile distilled water at a concentration of 10^8^ CFU⋅mL^−1^. Twenty μL of this bacterial suspension and the same volume of WGA solution were added to a slide glass and mixed well. Then the samples were air dried after 30 min moisturizing reaction at room temperature. The samples were stained with crystal violet, then were observed their agglutination reactions by a microscope. Phosphate-buffered saline instead of WGA solution was used as control. Isolates that were positive for WGA affinity were stored for further analysis.

Plant growth-promoting traits (34, 35), physiological and biochemical characteristics (36) of the bacterial isolates were examined. Meanwhile, these affinity isolates were identified based on 16S rRNA sequencing. One excellent affinity phosphate-solubilizing strain to wheat was selected for further studies.

### Electrotransformation of strain P34

*E. coli* WA803 strain was cultured in LB medium for about 15 h, and then the plasmid pTR102 was extracted using a SanPrep column plasmid DNA miniprep kit (Sangon Biotech Co., Ltd, ShangHai, China). One of the isolates, phosphate-solubilizing strain P34, was prepared to obtain competent cells for electrotransformation as follows: after overnight cultivation in LB medium at 28°C, cells of strain P34 were collected by centrifugation at 7000 g for 10 min at 4°C. Cells were washed with ice-cold ultrapure water, followed by washing with 10% (v/v) ice-cold glycerol, and then re-suspended in 50 μL 10% glycerol. The mixture of 5 μL plasmid and 50 μL competent cells was electroporated at 2.5 kV, 200 Ω, and 25 μF. The sample was re-suspended in 1 mL SOC medium and incubated at 28°C for 1.5 h with shaking at 160 r⋅min^−1^, and then spread on plates of LB medium containing 50 μg⋅mL^−1^ kanamycin. After culture for 2 or 3 days at 28°C, 10% capraldehyde was added in the dark and the colonies that fluoresced were selected as the transformants.

### Identification, genetic stability and plant growth-promoting traits of strain P34-L

Identification of strain P34-L was based on 16S rRNA gene sequencing. DNA was extracted and purified using a Sangon Bacterial Genomic DNA Extraction UNIQ-10 kit according to the manufacturer’s instructions. Amplification of the 16S rRNA gene sequences was carried out using primers 27F and 1492R (37). The PCR reactions were initiated with a denaturation step of 94°C for 4 min, followed by 30 cycles of denaturation at 94°C for 30 s, annealing at 55°C for 30 s, and extension at 72 °C for 60 s, with a final elongation step for 10 min at 72°C. PCR amplicons were submitted to Sangon Biotech (Shanghai) Co., Ltd for DNA sequencing. The resulting sequences were subjected to BLAST analysis with the NCBI database (http://blast.ncbi.nlm.nih.gov/Blast.cgi).

Cells of the transformant were plated successively on LB medium containing 50 μg⋅mL^−1^ kanamycin. Genetic stability of the transformant P34-L was judged based on whether the colonies have luminescence activity and kanamycin resistance.

Phosphate solubilization and siderophore production of strain P34-L was determined through the phospho-molybdate blue color method (34), and the Chrome Azurol S assay (35), respectively.

### Wheat seed pre-germination and planting

Wheat seeds were placed in sterile Petri dishes with water and kept in the dark at 25°C for about 2 days until their root length reached 2~3 mm. A liquid culture was prepared by culturing strain P34-L in LB medium for 15 h at 28°C with shaking (160 r⋅min^−1^) until the OD_600_ reached 1.0 (bacteria concentration at about 10^9^ CFU⋅mL^−1^). The germinated seeds were soaked in the bacterial solution for 30 min, and then sown in unsterilized yellow-cinnamon soil in rhizoboxes (one seed per rhizobox; 25 cm long × 13 cm wide × 1 cm thick). The plants were grown outdoors in natural surroundings.

### Wheat seedling harvest and treatment

Wheat seedlings were carefully removed from the rhizoboxes on days 6, 12, and 36 of growth to evaluate colonization and growth-promoting effects, and for qRT-PCR analyses on days 20, 30, 40, and 50. Soil particles were shaken off from the plant roots as carefully as possible before sampling to obtain whole roots. To evaluate colonization, wheat roots were cut into 2-cm long segments from the top to the bottom. Each root segment was placed in a flask with 100 mL sterilized water and was shaken vigorously for 1 h to release the attached cells. Serial dilutions were immediately prepared, and 0.1 mL of each dilution was spread on solid LB medium. The plates were incubated for 3 days at 28°C and the number of bacteria was determined by fluorescence observations. This value was used to calculate the number of P34-L colonizing the rhizosphere of each root segment. To analyze the effects of the bacterium on wheat plant growth, the wheat roots were rinsed in water, and then total root length (TRL), root projection area (RPA), root surface area (RSA), root volume (RV), root average diameter (RAD), and the number of root tips, forks, and crossings were measured using a WinRHIZO scanner (WinRHIZO Root Analysis, Regent Instruments, Quebec, Canada). At the same time, samples of wheat were cleaned with water for measuring their seedling fresh weight and root fresh weight, and then dried at 80°C to constant weight for measuring their seedling dry weight and root dry weight. Phosphorus contents in 30 d growth wheat leaves were measured by the vanadomolybdate blue method (38) after H_2_SO_4_-H_2_O_2_ digestion for leaves. Root samples were frozen in liquid nitrogen and stored at −80°C for qRT-PCR analyses. Data were subjected to one-way analysis of variance (ANOVA) with least significant difference (LSD) tests to detect significant differences between treatments and their controls. All statistical analyses were conducted using SPSS software (SPSS, Chicago, IL, USA). All the treatments were repeated 3 times, and data were average values of 3 replicates.

### RNA extraction and cDNA synthesis

Root samples were ground to a fine powder in liquid nitrogen, and RNA was extracted using an RNAiso plus kit (TaKaRa, Dalian, China). The RNA concentration was checked with a NanoDrop 2000 Spectrophotometer (Thermo Fisher Scientific Inc., Wilmington, DE, USA), and its quality was assessed by electrophoresis. First-strand cDNA was synthesized using the Prime Script™ RT reagent kit with gDNA Eraser (TaKaRa, Dalian, China) (39). The concentration of each RNA sample was adjusted to 1 μg per 20 μL reaction system. The cDNAs were diluted 10-fold in sterile distilled water before use in qRT-PCR analyses.

### Design and validation of primers

Several housekeeping genes have been tested as candidate reference genes for wheat. The housekeeping gene *Tubb* was used as the reference gene in this study for its high stability (primers: F, 5’-CAAGGAGGTGGACGAGCAGATG-3’ and R, 5’-GACTTGACGTTGTTGGGGATCCA -3’, amplicon length: 84 bp) (40). The *TaPT4* gene primers used were F (5’-GGGCTTCATGTTCACCTTCCTCGT -3’) and R (5’-GGATCTTTGACCCACTCAACATCC -3’) designed by Primer Premier 5.0 software, and the length of the amplicon obtained for this set of primers is 198 bp. Both sets of primers were validated by PCR under the following conditions: denaturation at 95°C for 5 min, and then 35 cycles of amplification at 95°C for 5 s, 58°C for 30 s, and 72°C for 20 s. The PCR products were tested by 1.2% agarose gel electrophoresis.

### Quantitative reverse-transcription PCR analyses

We used SYBR^®^ Premix Ex Taq™ II (TliRNaseH Plus) (TaKaRa) for qRT-PCR analyses (41). The qRT-PCR analyses were conducted using the Bio-Rad CFX96TM Real-Time System (Bio-Rad, Hercules, CA, USA). The PCR conditions were as follows: denaturation at 95°C for 30 s, followed by 40 cycles at 95°C for 5 s and 58°C for 30 s. Melting curve analyses were conducted before concluding the program with a 4°C hold. All samples were analyzed in triplicate. The relative transcript levels were calculated using the 2^−ΔΔCt^ method following the protocol of Livak and Schmittgen (42).

## Results

### Isolation of phosphate -solubilizing PGPR with WGA affinity

A total of 68 bacterial isolates were obtained from wheat rhizosphere based on a primary screening by phosphate growth medium, and 22 of them were determined positive for WGA affinity. Among all the tested WGA affinity isolates, the strain P34 was found to have excellent phosphate -solubilizing ability and plant growth-promoting traits, so the isolate, P34 was further assayed for rhizosphere colonization, wheat growth-promoting effects and phosphate transporter gene expression regulation as research material. Plant growth-promoting traits, physiological and biochemical characteristics of phosphate-solubilizing strain P34 are shown in Table 1. Strain P34 solubilized phosphate at 101.6 μg/mL, and produced siderophore at high amounts (+++++), which means the isolate exhibited a great ability in both of this two plant growth-promoting traits. The colonies of the isolate were regular and yellowish white in color, the cells were Gram-negative, rod-shaped, and non-endospore forming. The catalase test and hydrolyzed gelatin were positive, and the production of H_2_S, methyl red test and voges proskauer test were negative. The strain could utilize glucose to produce acid. The phosphate-solubilizing strain P34, which showed an affinity to WGA was further identified by 16 SrRNA sequence analysis (Fig.S1, for reviewers only), and ascertained as a *Pseudomonas* sp. (GenBank Accession number MF668124).

**Table 1.**
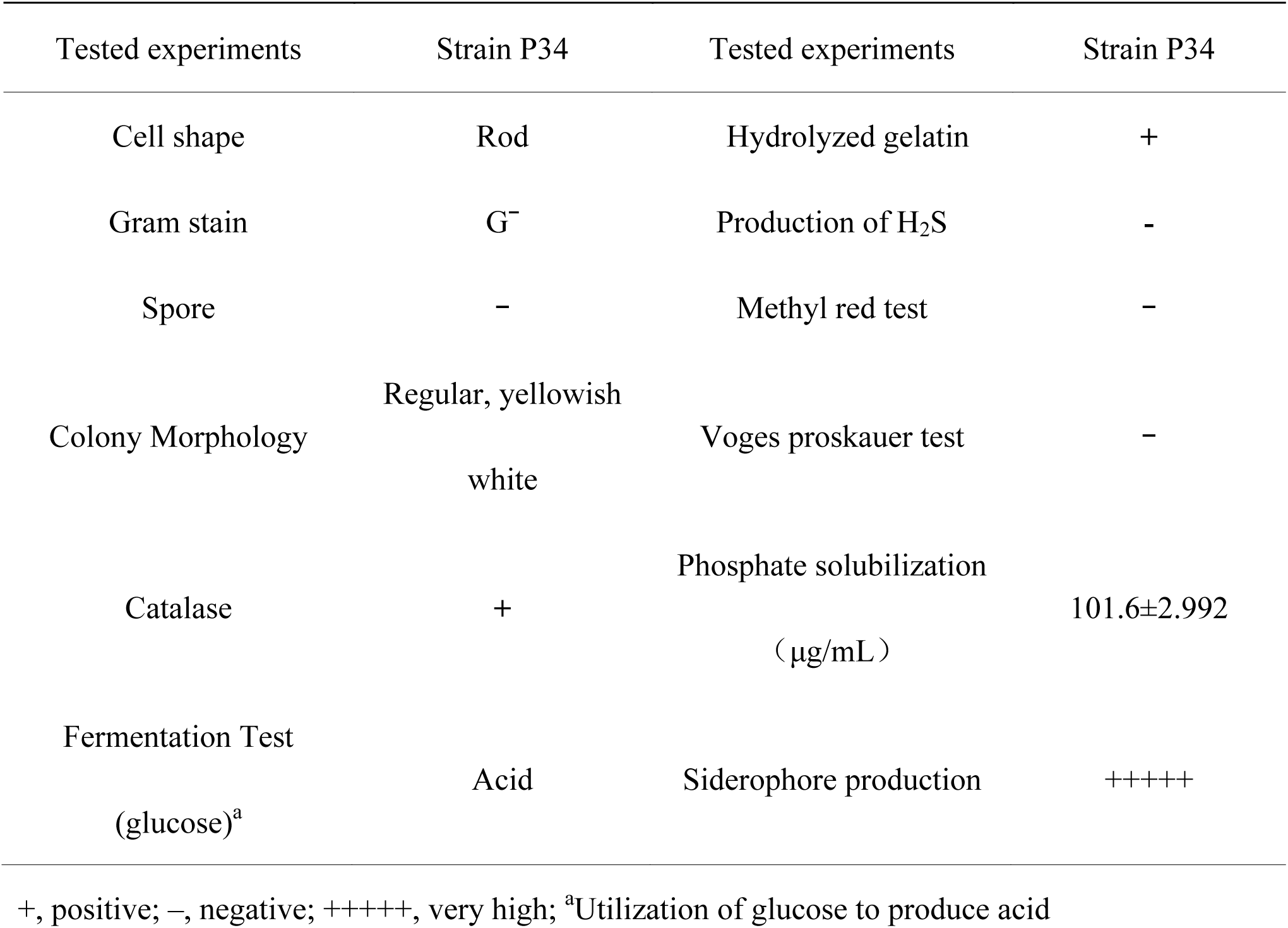
Characteristics of strain P34

### Screening, identification and genetic stability of the transformant strain P34-L

The *lux*AB gene was introduced into strain P34 by electrotransformation. Colonies of strain *Pseudomonas* sp. P34-L emitted fluorescence when 10% capraldehyde was added in the dark (Fig. 1). The nucleotide sequences of the 16S rDNA in P34-L is the same as those in P34 (GenBank Accession number MF668125). These results indicatied that *lux*AB was introduced successfully. Phosphate solubilization and siderophore production, two important characterizations of plant growth-promoting traits for bacteria have not appeared different from P34-L and P34 (Table S1, for reviewers only). After growth on plates of LB medium containing 50 μg⋅mL^−1^ kanamycin for 10 transfers, all colonies of P34-L retained luminescence activity and kanamycin resistance, indicating that P34-L was genetically stable.

**Fig. 1.**
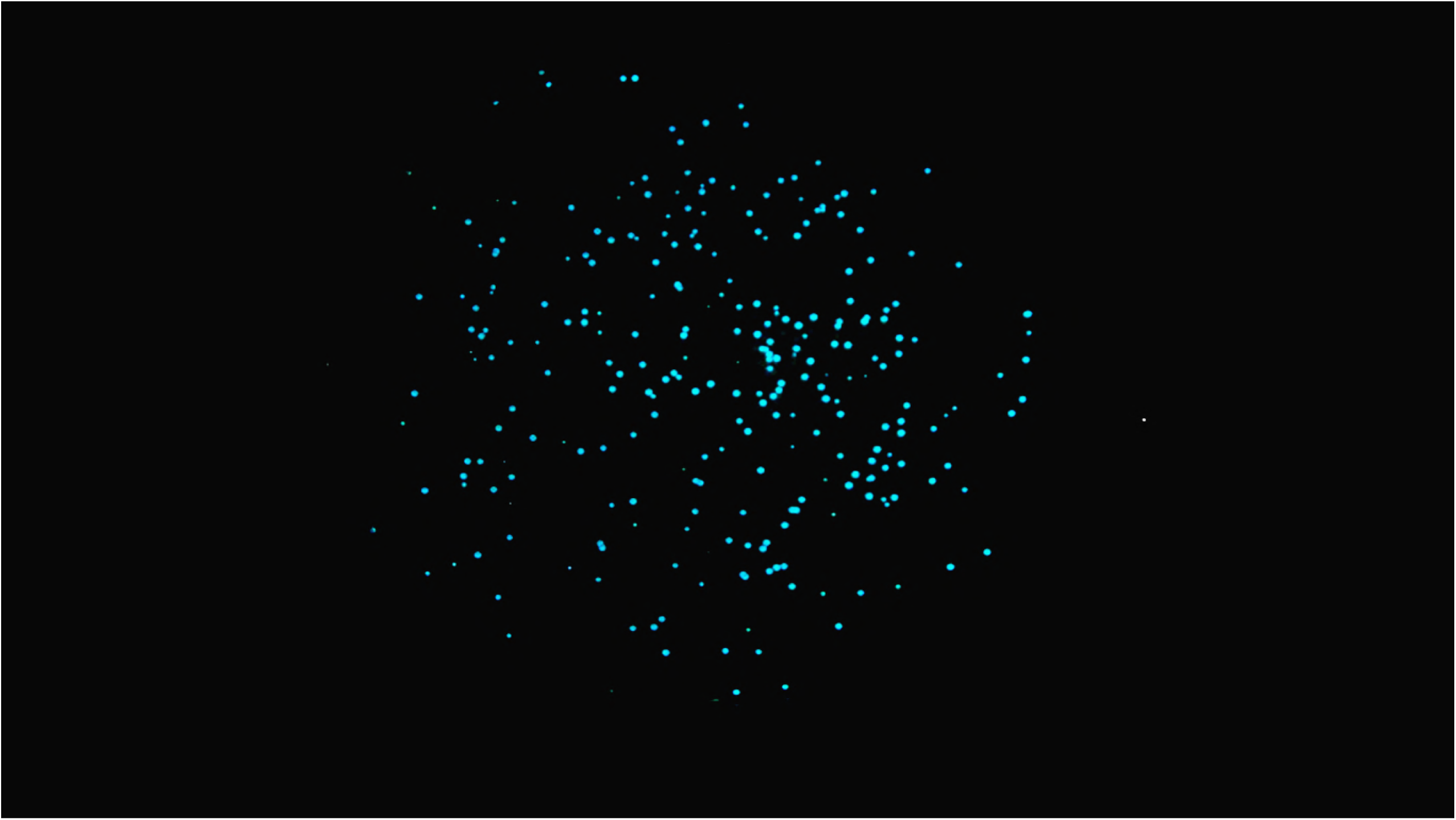
Fluorescence emitted by P34-L colonies

### Colonization dynamics and distribution of P34-L in the wheat rhizosphere

The labeled strain P34-L harboring the *lux*AB gene was inoculated on the surface of germinated wheat seeds for detecting the time and space characteristics of colonization. When the wheat seeds were planted, the labeled P34-L strain was present only the surface of germinated wheat roots about 0.3 cm long. By day 6, P34-L had spread to a depth of 8 cm below the wheat seeds, indicating that this strain was able to propagate and survive in new spaces as the roots grew. As indicated by the density of P34-L on days 12 and 36 of wheat growth, P34-L migrated from the surface to deeper layers as the roots grew. The colonization density of P34-L in all root segments of wheat decreased after 12 days of growth compared with that at 6 days, which may be due to fact that other indigenous microorganisms in the soil compete with the labeled strains for nutrition and favorable sites, thereby consuming nutrients in the soil. At the same time, the soil temperature, soil water content and other factors are also the reasons for the decrease of colonization density. On day 36, the density of P34-L at >8 cm root depth was 1.1×10^7^ cfu⋅g^−1^, higher than the density at 0~2 cm. The density of P34-L remained at a high level in deeper layers of the wheat rhizosphere (Table 2). As the roots grew, the original 0~2 cm root segment became an old root. The available root exudates for microbial growth decreased with the concentration of exudates secreted by old roots reducing. Consequently, the labeled strains adsorbed at 0 ~ 2 cm root segment gradually decreased. While P34-L moved to the young roots and colonized the rhizosphere of newly growing roots to became established at the radical. These results indicated that strain P34-L could survive in the wheat rhizosphere for a long time, and colonize new spaces in wheat rhizosphere following the extension of wheat roots. P34-L colonized the wheat rhizosphere successfully and stably, which provide the premise and basis for the strain to play the promoting growth role for wheat.

**Table 2.**
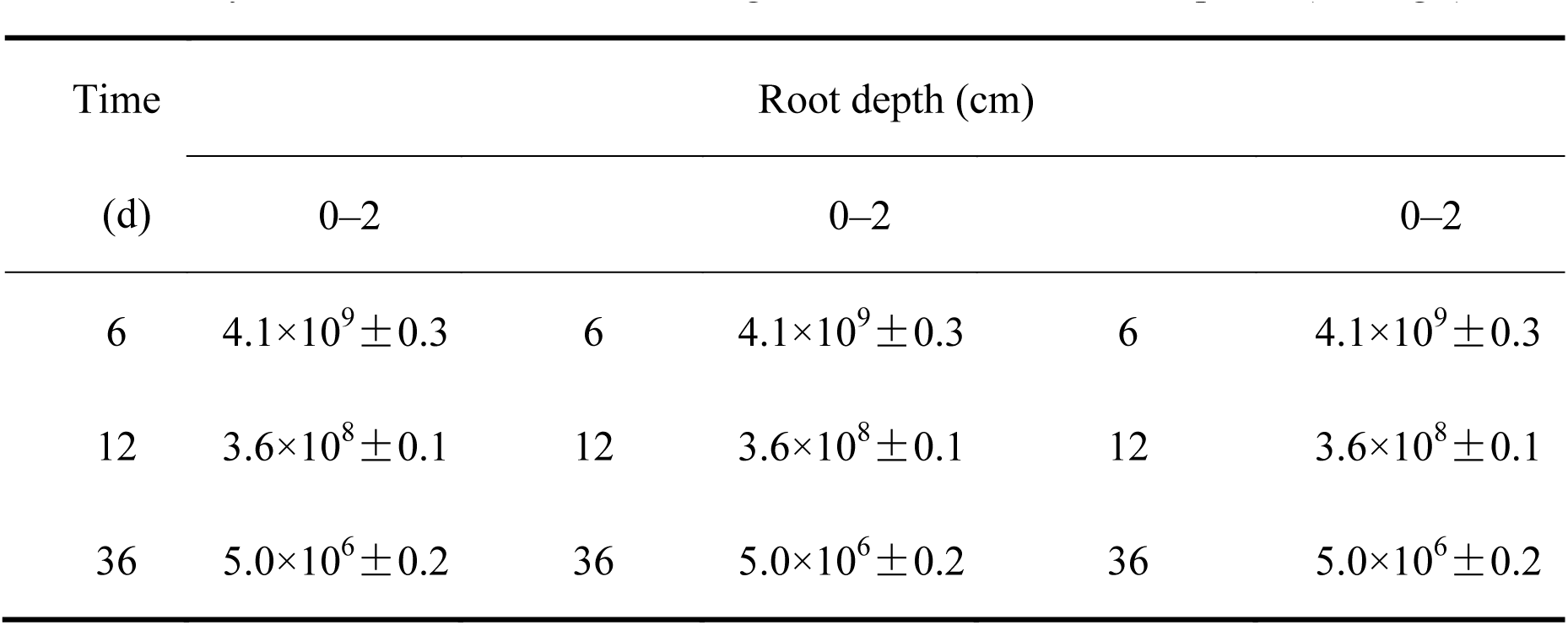
Density of P34-L on different root segments in the wheat rhizosphere (CFU⋅g^−1^)

### Growth-promoting effects of strain P34-L

To investigate the effects of strain P34-L on the growth of wheat, we measured the seedling fresh weight, seedling dry weight, root fresh weight and root dry weight of wheat plants at different growth stages (Table 3). At the beginning of the wheat growth period, there was no difference between uninoculated and inoculated wheat plants. By day 36, the seedling fresh weight, seedling dry weight, root fresh weight, and root dry weight of inoculated plants were 100.73%, 94.42%, 50.84%, and 31.85% higher, respectively, than those of control plants (p<0.01). This result indicated that strain P34-L remarkedly promoted dry matter accumulation in wheat plants.

**Table 3.**
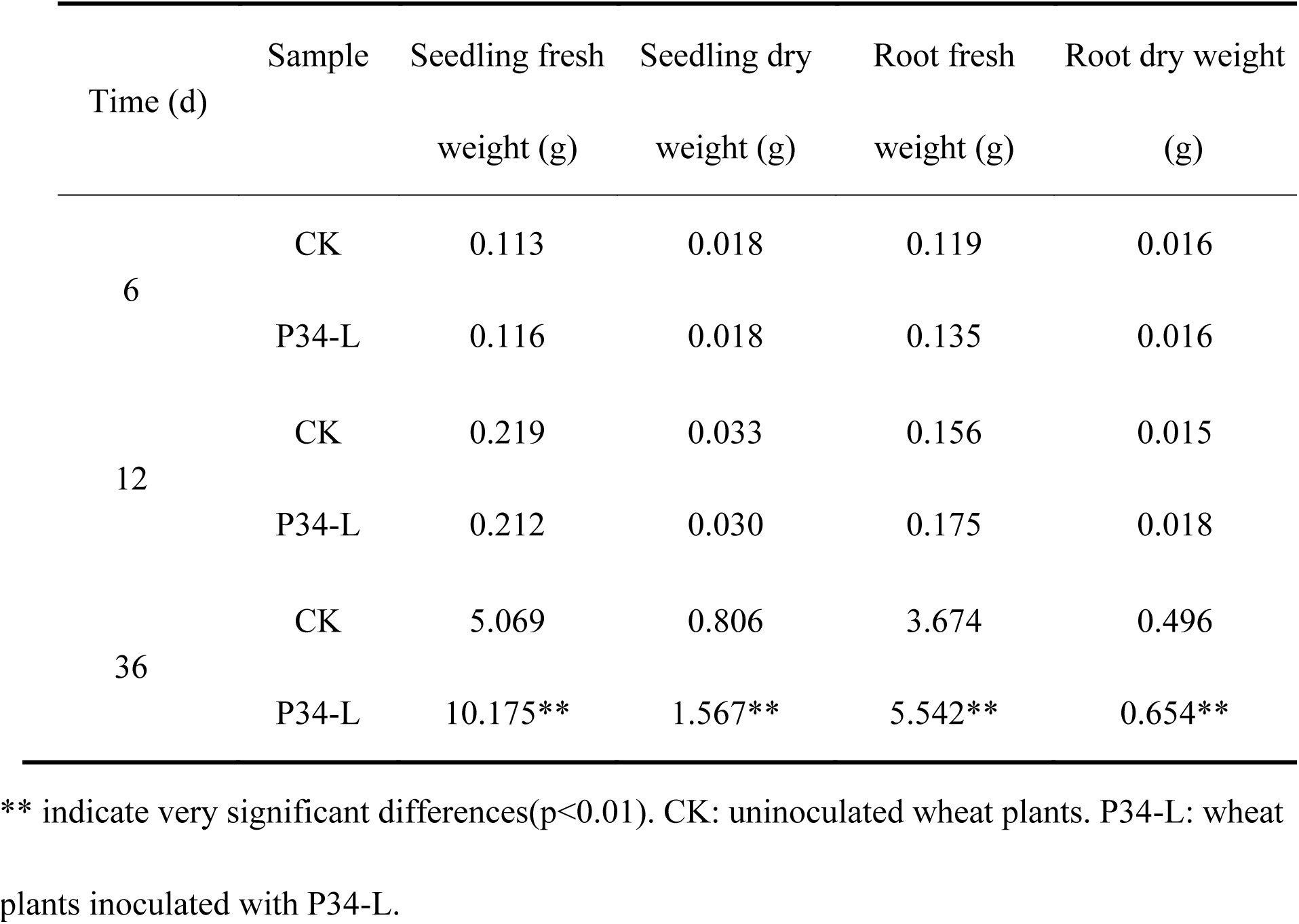
Effects of strain P34-L on fresh matter and dry matter accumulation of wheat

The total root length (TRL), root projection area (RPA), root surface area (RSA), root average diameter (RAD), root volume (RV), and the number of root tips, forks, and crossings of wheat were measured using a WinRHIZO scanner (Table 4). By day 12, P34-L had a certain promotion effect on wheat root growth. By day 36, the TRL, and number of tips, forks, and crossings were 95.12%, 184.16%, 78.86%, and 132.14% higher, respectively, in inoculated wheat plants than in control ones (p<0.01). Root systems morphology of 30 d growth wheat plants were shown in Fig.2. From the figure it is clear that wheat plants inoculated with P34-L have more lateral roots and lager root systems compared to those uninoculated ones. Strain P34-L promoted the development of wheat root system. Developed roots make it easier for wheat to absorb nutrients and water, which is beneficial to the growth of wheat.

**Fig. 2.**
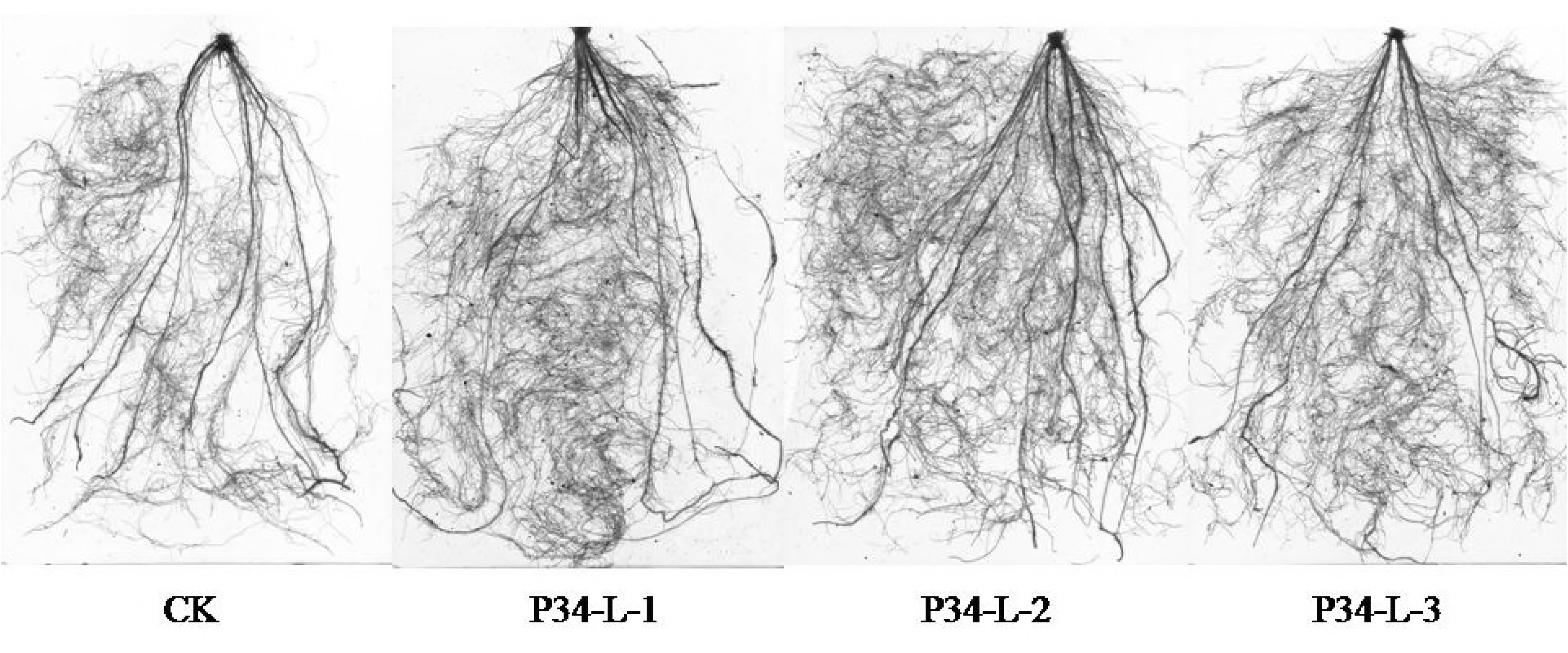
Root systems morphology of 30 d growth wheat plants. CK: root systems for uninoculated wheat plants. P34-L-1, P34-L-2, P34-L-3: root systems for wheat plants inoculated with P34-L.

**Table 4.** Effects of strain P34-L on wheat root growth

| Time<br>(d) | Sample | TRL<br>(cm) | RPA<br>(cm<sup>2</sup>) | RSA<br>(cm<sup>2</sup>) | RAD<br>(mm) | RV<br>(cm<sup>3</sup>) | Tips | Forks | Crossings |
| --- | --- | --- | --- | --- | --- | --- | --- | --- | --- |
| 6 | CK | 203.30 | 7.72 | 24.26 | 0.38 | 0.23 | 371.50 | 1277.50 | 19.00 |
|  | P34-L | 222.50 | 7.69 | 24.15 | 0.35 | 0.21 | 589.50 | 1247.00 | 19.00 |
| 12 | CK | 208.20 | 6.89 | 21.65 | 0.33 | 0.18 | 280.50 | 655.50 | 110.00 |
|  | P34-L | 240.00 | 7.76 | 24.38 | 0.32 | 0.20 | 305.50 | 855.00** | 132.50 |
| 36 | CK | 2121.00 | 164.00 | 515.21 | 0.78 | 9.99 | 6143.50 | 26454.00 | 3336.50 |
|  | P34-L | 4138.50* | 204.40 | 641.98 | 0.49* | 7.93 | 17457.50** | 47316.50* | 7745.50** |

The phosphorus content of the 30th day dried leaves was measured (Fig.3). The phosphorus content of leaves in inoculated wheat reached 0.49 mg⋅g^−1^, which was 25.64% higher than in uninoculated wheat (p<0.01), which means phosphate -solubilizing strain P34-L can enhance phosphorus accumulation in wheat leaves effectively.

**Fig. 3.**
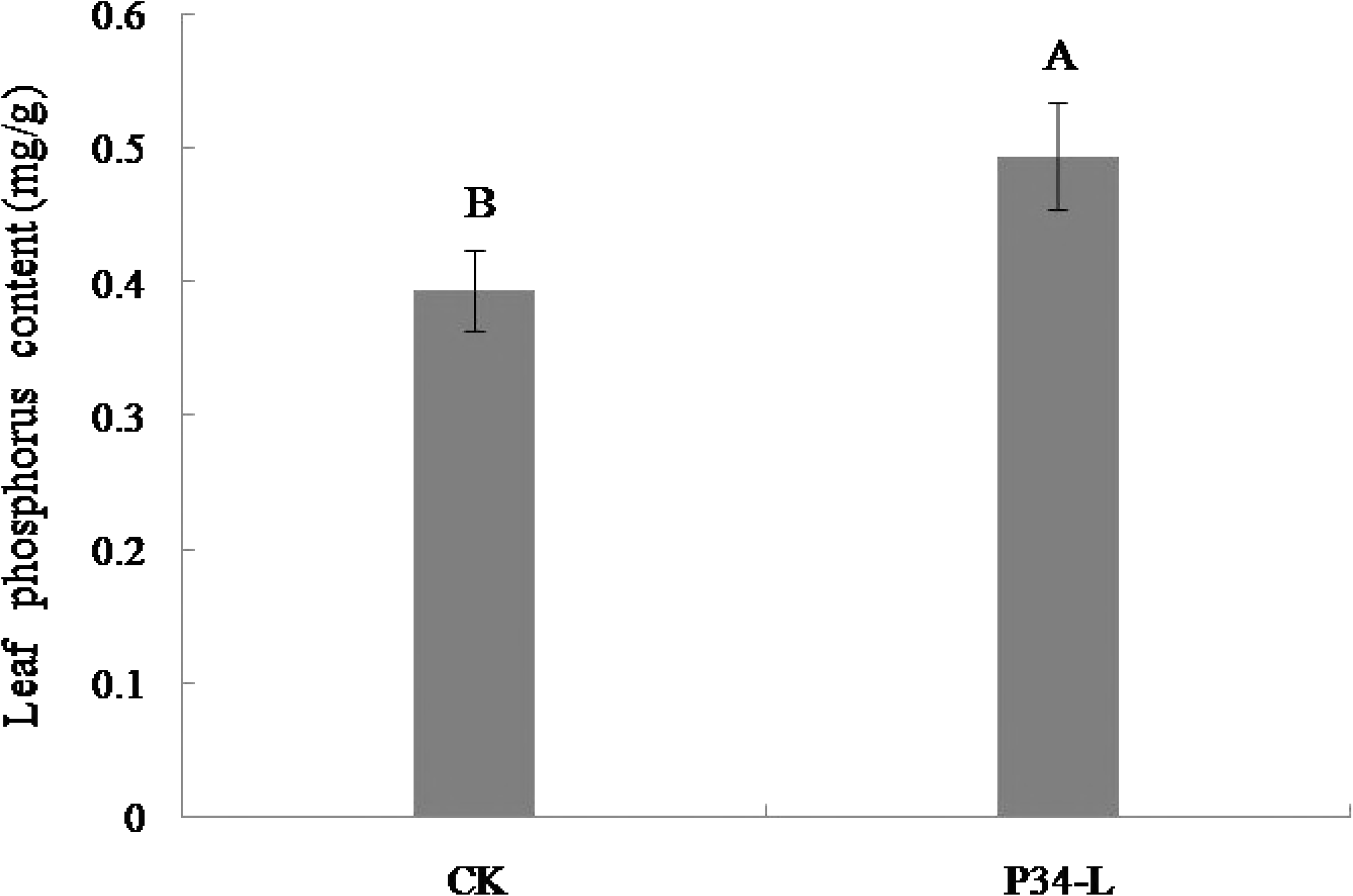
Effect of p34-L on phosphorus accumulation in 30th day growth wheat leaves. Data are average values of three independent experiments ± SE. CK: uninoculated wheat plants. P34-L: wheat plants inoculated with P34-L.

Compared with uninoculated wheat plants, those inoculated with P34-L showed significantly increased phosphorus accumulation in leaves, above-ground fresh weight, above-ground dry weight, root fresh weight, root dry weight, total root length, and number of root tips, forks, and crossings. The results indicated that P34-L is of significant value for wheat production improvement by promoting the root growth and dry matter accumulation. Strain P34-L may be a potential ideal biofertilizer producing strain for sustainable agriculture.

### Relative transcript levels of phosphate transporter gene *TaPT4* in wheat roots

To verify the phosphate-solubilizing ability of strain P34-L in gene level and investigate its molecular mechanism of phosphorus solubilizing as well, the transcript level of a phosphate starvation-inducible phosphate transporter gene *TaPT4* in wheat roots was monitored by quantitative reverse-transcription PCR. The results were shown in Fig.4. The relative transcript levels of *TaPT4* were lower in inoculated wheat plants than in uninoculated ones after 20-50 days of wheat growth. The relative transcript level of *TaPT4* were 52.46%, 53.47%, 83.50% and 83.00% lower in inoculated plants of 20d, 30d, 40d, 50d growth than in control plants, respectively (p<0.01). This result indicated that strain P34-L down-regulated the transcript level of *TaPT4* in wheat roots, which means a better phosphorus nutritional status have been established in the rhizosphere of wheat by the strain. The outcome of the experiment provides effective evidence for strain P34-L in possessing good ability to dissolve phosphorus in gene level.

**Fig. 4.**
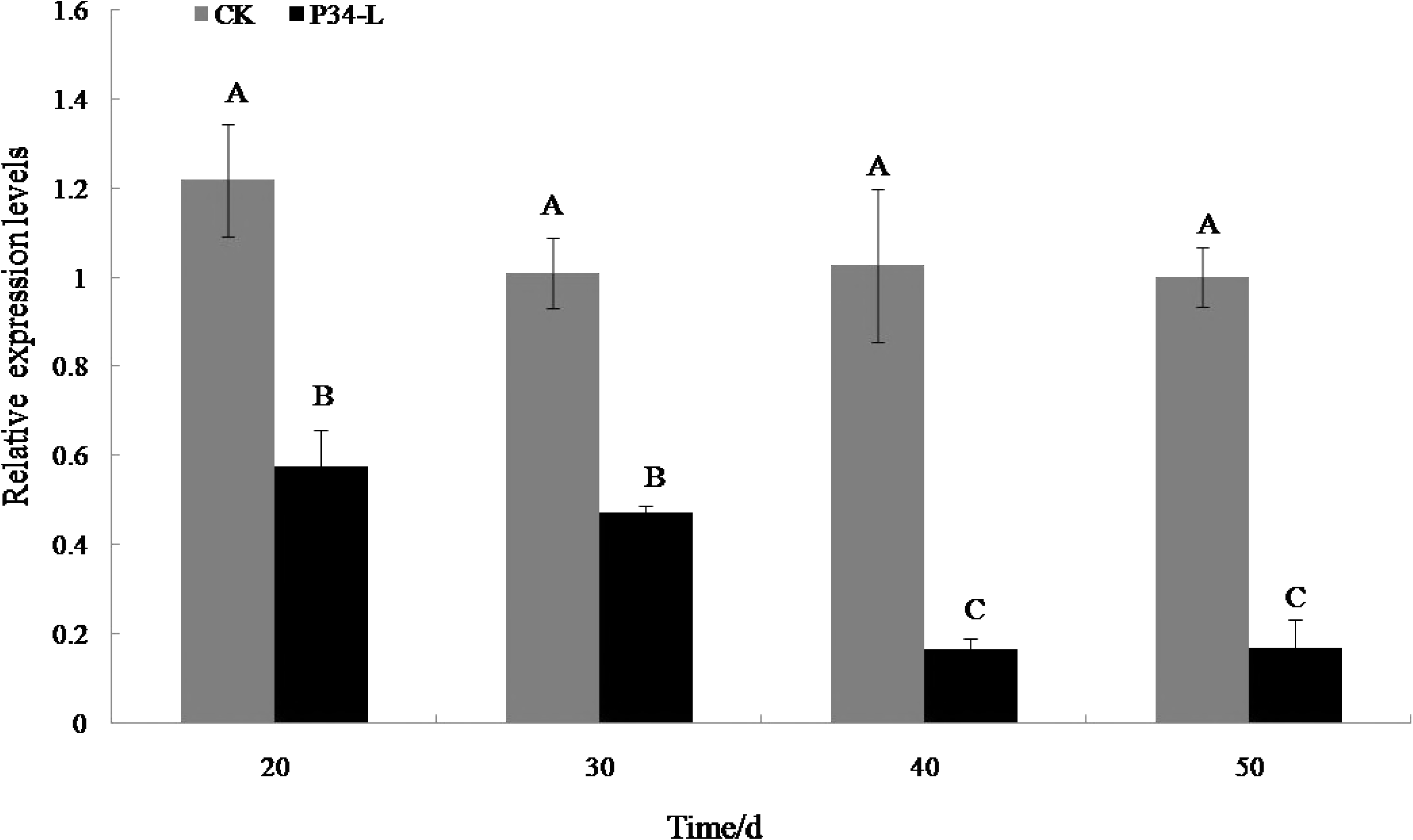
Relative transcript levels of phosphate transporter gene *TaPT4* in wheat roots. Data are average values of three independent experiments ± SE. CK: uninoculated wheat plants. P34-L: wheat plants inoculated with P34-L.

## Discussion and conclusions

### Colonization of the wheat rhizosphere by P34-L and its growth-promoting effects on wheat

Beneficial interactions between plants and microbes in the rhizosphere are a vital factor for plant development and health (43, 44). PGPR benefit by utilization of plant roots exudates as a source of nutrients for their growth and promote the growth of their host plant in several ways, including fixing atmospheric nitrogen, solubilizing phosphorus and potassium, producing phytohormones, and suppressing plant pathogens (44–48). In spite of the importance of PGPR for plant growth, few studies have focused on the crucial steps of how these microbes stably colonize the plant rhizosphere. Nevertheless, the stable and successful colonization of PGPR in the rhizosphere of the host plant is a key step for a successful PGPR–host relationship. The ability to successfully colonize the rhizosphere is an important basics requirement for field application of PGPR strains.

There is a complex process of signal exchange and mutual recognition between microbes and their plant host. The precursor materials identified so far are lectins secreted by plants and extracellular polysaccharides secreted by bacteria (11–13, 49–51). Although the relationship between PGPR and plants is weaker than that between rhizobia and legumes, there is a certain affinity between them. To explore this relationship, a total of 22 phosphate-solubilizing strains were obtained from the wheat rhizosphere by a new screening method of selective medium combined with WGA in this study. These isolates showed an affinity to wheat germ agglutinin. Among which we focused on an excellent phosphate-solubilizing *Pseudomonas* sp. strain P34 for study its colonization in wheat rhizosphere, effects on wheat growth, and the expression level of phosphate transporter gene *TaPT4* in wheat root.

The luminescent enzyme gene marker is a modern technique for tracing the colonization dynamics of PGPR in plant rhizosphere (52). The ineffectiveness of PGPR for plant growth promotion has been attributed to the inability of the strain to colonize the plant rhizophere (53). Therefore, the ability to successfully colonize the root is an important basics requirement for field application of strains. Arikan and Pirlak reported that application of sour cherry growth promoting rhizobacteria T8 and OSU-142 significantly increased the yield, leaf area and shoot length of sour cherry as compared to the control (54). Bal et al. obtained three ACC deaminase producing PGPR isolates from rice rhizosphere, and found these selected bacterial isolates had considerable positive impacts on growth parameters of rice including root length, shoot length and weight, germination percentage and chlorophyll content, compared with the uninoculated control (55). Mirzaei et al. studied the effects of PGPR (Azotobacter) on growth promoting characteristics and colonization efficiency in fennel. The results showed that Azotobacter can significantly increase the root colonization percentage, grain yield and biological yield of fennel (56). These studies focused on the promoting growth effects and the colonization level of PGPR in the rhizosphere of crops, but did not link the colonization ability of the strain with the development of the root system. However, the effective colonization of the strain not only means that the strain can survive in the rhizosphere of the plant, but also can grow and propagate on the new root with the elongation of the root. In our work, we investigated the temporal and spatial characteristics of colonization of PGPR in wheat rhizosphere and its effects on wheat growth, root development, and phosphorus accumulation in wheat. To make tracing easier, the plasmid pTR102 harboring the luciferase *lux*AB gene was transferred into a phosphate-solubilizing *Pseudomonas* sp. strain to create P34-L. The labeled strain was then used as research material in this study. The results indicated that strain P34-L has good affinity with wheat. It can colonize the wheat rhizosphere for a long time, and can continue to colonize with the elongation of the root. Moreover, the labeled stain can promote the dry matter accumulation, increase total root length, number of root tips, forks, and crossings of wheat, and enhance phosphorus accumulation in wheat leaves. It is an indication that it increased phosphorus accumulation in wheat plants, promoted the development of the wheat root system and can play a positive role in the growth and development of wheat. Strain P34-L can survive in wheat rhizosphere in a long time, indicating that the strain has strong rhizosphere colonization ability and high rhizosphere environmental adaptability. The strain with colonization ability can live and propagate in the rhizosphere of wheat for a long time, and continuously produce favorable substances for plant growth. It can make full use of root exudates to promote the root development of wheat, and then directly affect the nutritional status of the above ground part, and promote the overall development of wheat. The ability of colonization is an important prerequisite and basis for studying the growth and development of wheat promoted by PGPR.

### Expression of phosphate transporter gene *TaPT4* in wheat in response to strain P34-L

The uptake and transportation of phosphorus in plants are mediated by phosphate transporters. They play pivotal roles in regulation of absorption and utilization of phosphorus for plants under diversified phosphate-supply conditions (57). As one member of high affinity phosphate transporter gene family, *TaPT4* was reported to play an important role in enhancing plant resistance to low phosphorus stress through enhanced expression (26, 57). Furthermore, *TaPT4* is also a phosphorus deficiency indicator gene. The transcription of *TaPT4* gene in roots was induced by low phosphorus stress, and down-regulated by phosphorus supplementation in wheat roots, so it was applied to reflect the phosphate level in environment (26). Tian et al. reported that the expression of *TaPT4* gene was down-regulated by arbuscular mycorrhizal fungi colonization when roots of wheat received phosphorus signals (26). In this work, we evaluated the effect of strain P34-L on the transcript level of *TaPT4* in wheat at different developmental stages. The relative transcript levels of *TaPT4* were lower in inoculated wheat roots than in uninoculated ones after 20-50 days of wheat growth. These results suggest that strain P34-L may solubilize insoluble phosphorus in soil into soluble phosphorus for plant absorption and utilization, increase the phosphorus content in the environment, and then down-regulated the transcript levels of *TaPT4* in wheat roots. One of the mechanisms by which P34-L promotes wheat growth might be that the strain releases phosphorus from the soil, provide sufficient phosphorus for the growth of wheat. At gene level, inoculation of strain P34-L can down-regulate the transcript levels of phosphate transporters *TaPT4* gene in wheat roots. These results lay a firm foundation for further research on the relationships between PGPR and their host plants. Furthermore, the results of our present work suggested that, phosphate-solubilizing *Pseudomonas* sp. strain P34 can colonize wheat rhizosphere, and have a great potential to increase the growth of wheat. Therefore, it may be utilized as bio-fertilizer for wheat production in sustainable agriculture systems.

## Acknowledgements

This work was supported by the National Natural Science Funds of China (41401269), the Key Projects for Exceptional Young Teachers in Anhui Province (2013SQR015ZD) and the Funds of Anhui Agricultural University (2013(5), wd2012-6).

